# Genome-Wide association study uncovers pea candidate genes and pathways involved in rust resistance

**DOI:** 10.1101/2024.04.10.588888

**Authors:** Salvador Osuna-Caballero, Diego Rubiales, Nicolas Rispail

**Author notes:** Corresponding author: Salvador Osuna-Caballero.

## Abstract

Pea is an important temperate legume crop providing plant-based proteins for food and feed worldwide. Pea yield can be limited by a number of biotic stresses, among which, rust represents a major limiting factor. Some efforts have been made to assess the natural variation in pea resistance, but its efficient exploitation in breeding is limited since the resistance loci identified so far are scarce and their responsible gene(s) unknown. To overcome this knowledge gap, a comprehensive Genome-Wide Association Study (GWAS) on pea rust, caused by *Uromyces pisi*, has been performed to uncover genetic loci associated with resistance. Utilizing a diverse collection of 320 pea accessions, we evaluated phenotypic responses to two rust isolates using both traditional methods and advanced image-based phenotyping. We detected 95 significant trait-marker associations using a set of 26,045 DArT-seq polymorphic markers. Our *in-silico* analysis identified 62 candidate genes putatively involved in rust resistance, grouped into different functional categories such as gene expression regulation, vesicle trafficking, cell wall biosynthesis, and hormonal signalling. This research highlights the potential of GWAS to identify resistance sources, molecular markers associated with resistance and candidate genes against pea rust, offering new targets for precision breeding. By integrating our findings with current breeding programs, we can facilitate the development of pea varieties with improved resistance to rust, contributing to sustainable agricultural practices and food security. This study sets the stage for future functional genomic analyses and the application of genomic selection approaches to enhance disease resistance in peas.

**Key message:** Candidate genes and metabolic pathways controlling resistance to rust disease in pea have been proposed through GWAS using 26,045 DArTseq polymorphic markers and phenotypic data from field and controlled conditions.

## 1 Introduction

Pea (*Pisum sativum* L. 2n = 14) is one of the principal legume crops grown globally, ranking as the third most produced pulse in the world after dry beans (FAOSTAT 2022). This crop is a vital source of plant-based protein for both animal feed and human food, offering significant health benefits (Tulbek et al. 2016; Clemente and Olias 2017). Additionally, pea plays a crucial role in agricultural sustainability through nitrogen fixation enhancing soil fertility and structure, making it a valuable component of crop rotation or intercropping systems, particularly in conjunction with cereals (Skoufogianni et al. 2019; Gungaabayar et al. 2023).

Peas are prone to various diseases and pests, which are principal factors limiting their yield (Rubiales et al. 2015, 2023). Rust disease poses a substantial threat in many pea-growing regions worldwide. Caused by two biotrophic fungi, *Uromyces viciae-fabae* (Pers. de Bary) in warm, humid climates, and *U. pisi* (Pers.) (Wint.) in temperate zones, rust can lead to significant yield reductions—up to 50% for *U. viciae-fabae* and 30% for *U. pisi* infections (Barilli et al. 2012a; Singh et al. 2023). Such impacts on yield and seed quality underscore the necessity for focused breeding and management strategies to effectively counteract rust disease in peas..

Rust control with fungicides is effective (Emeran et al. 2011) but requires repeated treatments which is hardly economical for low input field crops like pea. Also, concerns are raised on environmental issues of pesticide control. To address this, efforts have been made to control pea rust inducing systemic acquired resistance (Barilli et al. 2009b) or using natural compounds, that were shown promising although the available formulations are only partly effective, and their long-term environmental impacts remain unverified (Barilli et al. 2012b, 2017, 2022). Therefore, further emphasis is needed to develop cultivars resistant to rust. In this line, large pea germplasm collections have been evaluated for resistance to *U. pisi*, disclosing and characterizing variable levels of incomplete resistance, with no complete resistance available so far (Barilli et al. 2009a; Osuna-Caballero et al. 2022). Partial resistance (PR) is characterized by a non-hypersensitive response and reduced disease severity (DS). It is very frequent in other legumes against various rusts, and it is the common source of resistance found in peas against rust (Rubiales et al. 2011). Conversely, hypersensitive resistance (HR), often monogenic in origin, has also been observed in other rust pathosystems caused by *U. viciae-fabae* in lentils and faba beans (Avila et al. 2003; Negussie et al. 2012; Rubiales et al. 2013; Adhikari et al. 2021). Recent screenings of a pea core collection have uncovered new sources of PR and identified for the first time a pea accession combining HR and PR against *U. pisi* (Osuna-Caballero et al. 2022). Furthermore, new, and more efficient image-based phenotyping methods are improving the limitations of traditional breeding in this pathosystem (Olivoto et al. 2022; Osuna-Caballero et al. 2023). Nevertheless, despite previous efforts to unravel the genetic control underlying resistance to this fungal disease—where QTL responsible for resistance in *P. fulvum* were identified—clarifying the genetic architecture of HR and PR to rust caused by *U. pisi* remains necessary (Barilli et al. 2018). Furthermore, identification of candidate genes could be instrumental in foreground selection of favourable alleles.

The phenotypic information available in pea germplasm, combined with new high-throughput phenotyping systems focused on rust (Gallego-Sánchez et al. 2020; Osuna-Caballero et al. 2023), and the availability of high-quality reference pea genomes (Kreplak et al. 2019; Yang et al. 2022), enables more in-depth genomic studies (Pandey et al. 2021). Genome-wide association studies (GWAS) have emerged as a powerful approach to dissect the complex genetic underpinnings of quantitative traits within diverse plant populations. By scanning the genome for polymorphisms correlated with phenotypic variance, GWAS has facilitated the identification of both single nucleotide polymorphisms (SNPs) and DArT-seq derived markers that are intricately linked with key traits, providing candidate genes associated with traits of interest across numerous legume species (Susmitha et al. 2023). Specifically, recent GWAS in peas have been useful to identify favourable alleles for agronomic traits or disease resistance (Alemu et al. 2022; Leprévost et al. 2023), but there is still a lack of such studies for rust disease.

In the present study, we bring new sources of resistance against two *U. pisi* isolates by employing both traditional screening methods and innovative image-based methodologies across an extensive germplasm collection of 320 pea accessions, representative of global diversity. Moreover, we explored the genetic architecture of partial resistance to rust infection through GWAS, as a preliminary step towards more efficient precision breeding. To achieve this, the phenotypic response of the world collection of pea accessions inoculated with *U. pisi* was integrated with previously generated DArT-seq polymorphic markers, revealing significant associations between them. Therefore, our in-silico analysis of the genes containing or in the vicinity of the associated markers led to the discovery of previously uncharted metabolic pathways potentially implicated in rust response.

## 2 Material and Methods

### 2.1 Plant and fungal material

Plant material consisted in a diverse core collection of 320 *Pisum* accessions of worldwide origin gathered previously at Institute for Sustainable Agriculture – CSIC to ensure an extensive range of characteristics in terms of phenotype and their genetic makeup (Rispail et al. 2023). The collection includes accessions that have already demonstrated their potential for resistance to several pea diseases including rust (accessions IFPI3260, PI347352, PI347343, PI347353 and PI347389) (Barilli et al. 2009a).

To assess the resistance levels within the collection to rust disease, we used in this study two highly virulent isolates of *Uromyces pisi* that specifically affect pea crop, UpCo-01 and UpKeS-05, preserved at IAS – CSIC (Barilli et al. 2012a).

### 2.2 Phenotyping

Three series of experiments were conducted to assess the rust resistance components of the pea collection. One series of experiments took place under field conditions over three cropping seasons, involving adult plants, while the other two experiments were conducted in controlled conditions (CC) using seedlings infected with the *U. pisi* isolate UpCo-01 or UpKeS-05.

For the field experiment, the pea collection was evaluated over three consecutive crop seasons from 2018 to 2020 in Cordoba, Spain, as described in Osuna-Caballero et al. (2022). Briefly, ten seeds per accession were sown in 1-meter rows, spaced 0.7 meters apart, in an alpha lattice design with three replicates. Mature plants were artificially inoculated with the *U. pisi* isolate UpCo-01, and Disease Severity (DS) was assessed as the percentage of rust coverage 30 days after inoculation (dai).

The first CC experiment, involved evaluating the pea collection against the UpCo-01 isolate, followed a randomized complete block design (RCBD), and the three inoculations involving two plants per genotype were performed on pea seedlings as described in Osuna-Caballero et al. (2022). Therefore, screening was done with six replicates per accession, visually counting pustules (Infection Frequency, IF) from 7 to 13 dai allowed to estimate the Area Under Disease Progress Curve (AUDPC), the Latency Period (LP_50_) and the Monocyclic Disease Progress rate (MDPr). Additionally, at 13 dai, DS and infection type (IT) was also visually evaluated.

The second CC experiment utilized the UpKeS-05 rust isolate and included an additional biological replicate in the three inoculations, resulting in a total of nine replicates per genotype. The inoculation procedure was identical to the previous CC experiment, ensuring consistency in how the plants were exposed to the rust. However, at 7 dai, when the rust colonies were already stablished, a detached leaflet from each plant was placed on a Petri dish with an agar-water mixture (1:10 v:v). This method, detailed in Barilli et al. (2022), was devised to preserve leaf viability, facilitating the documentation of rust progression via imaging. Subsequent analysis of daily photographs of these Petri dishes, spanning from 7 to 13 dai, was conducted using the *pliman* package in R software (Olivoto et al. 2022). This analysis aimed to quantify infection frequency (IF), disease severity (DS), and pustule size (PS, mm^2^), in addition to calculating the corresponding disease progression parameters (AUDPC, LP_50_, and MDPr) - except for PS. The use of the RGB evaluation method, as described in Osuna-Caballero et al. (2023) underpins this approach, which has been shown to achieve a high level of concordance with traditional visual evaluations. By maintaining consistency in the inoculation procedures across both CC experiments and employing an image-based evaluation method with proven reliability and comparability to visual assessments, we assert that the results from these experiments are indeed comparable.

### 2.3 Phenotypic data analysis

The statistical analysis of the three phenotypic data sets (field, CC_UpCo-01_, and CC_UpKeS-05_) was conducted using R 4.2.2 (R Core Team, 2021) and the *lme4* package (Bates et al. 2015) for mixed models. The linear model with interaction effect used to analyse the field data from multi-environment trials was applied using the genotype effect as fixed factor, while the remaining sources of variance, such as environment, environment x genotype interaction, and replicated block nested in environments, were included as random effects variables. Prior to analysis, field data set underwent an arcsine transformation to achieve normal distribution of disease severity percentages (DS), which was confirmed by checking the assumptions of residual normality and variance homogeneity. Heritability on the mean basis (H^2^) was calculated following the formula from Olivoto and Lucio (2020):

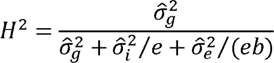

Where 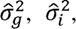 and 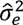 are the genotypic, the genotype-by-environment interaction, and the residual variance, respectively; and e and b are the number of environments and blocks, respectively.

For both controlled conditions (CC) phenotypic data sets, DS was transformed with an arcsine transformation, and other parameters related to disease progression and daily resistant components (IF and PS) were transformed with a square root transformation to achieve normal distribution. The genotype effect was included as a fixed factor, and the replicate, block, and combination of genotype and replicate were treated as random effects variables in the mixed model analysis. In case of CC, the H^2^ was calculated as follows:

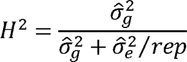

Where rep means the number of replicates within the experiment (Olivoto and Lucio 2020).

For the three data sets, mean separation between accessions and the overall mean was determined using Dunnett’s test (*p* ≤□0.05). The distributions are presented as best linear unbiased estimations (BLUEs) obtained from the mixed model. Correlation between estimated means under controlled or field conditions was assessed using the Pearson correlation coefficient. To simplify the complexity of the 19 traits obtained, a principal component analysis (PCA) was performed using *factoextra* R package (Kassambara and Mundt 2020).

### 2.4 Genotyping

The process of DNA extraction, DArTseq protocol, and data analysis were previously detailed in Rispail et al. (2023). In summary, the genotyping of the pea core collection was carried out using the DArTSeq approach, as described by Barilli et al. (2018). DNA was extracted from pooled leaves of 20 seedlings per accession and subjected to high-density Pea DArTSeq 1.0 array analysis. Data cleaning was performed to remove markers with low quality or lacking polymorphism, following the recommendations of Pavan et al. (2020). As a result, 26,045 polymorphic Silico-DArT markers were retrieved. The filtered markers were then mapped onto the improved *Pisum* reference genome sequences (Kreplak et al. 2019; Yang et al. 2022).

### 2.5 Genome-wide association mapping

The 26,045 genome-wide DArTseq markers were used for association mapping of rust in the GWAS panel of 320 pea accessions. In this study, a single-trait GWAS was conducted using two different models: the single locus mixed linear model (MLM) and the multi-locus Bayesian information and linkage disequilibrium iteratively nested keyway (BLINK) model (Huang et al. 2019). Both models were performed within GAPIT 3.0 (Wang and Zhang 2021) and have different assumptions: the MLM model allows for the identification of individual markers associated with the trait, while the BLINK model considers multiple markers simultaneously, considering the joint effects of nearby markers. Therefore, the analysis can capture both major and minor genetic effects contributing to the rust response variation.

The Bayesian information criterion (BIC)-based model selection procedure was used to determine the optimum number of principal components (PCs) required to efficiently control for population structure of the collection. Additionally, the Astle kinship matrix was used to estimate relatedness among individuals, further accounting for potential genetic correlations (Astle and Balding 2009).

To determine significant DArTseq markers, two different threshold levels were applied. The Bonferroni corrected LOD threshold, calculated based on the number of markers (26,045), offers a more stringent criterion to minimize false positives. On the other hand, the false discovery rate (FDR) method, proposed by Storey and Tibshirani (2003), provides a less restrictive threshold, allowing for the identification of potentially interesting markers that might be missed by the Bonferroni correction. FDR limit was set for each model and trait combination to ensure less than one false positive association with the R package *qvalue* (Storey and Bass 2022).

To visualize the results, Manhattan plots were generated, displaying the -log10(p-value) for each marker along the seven chromosomes, highlighting regions where significant associations occur. Quantile-quantile (Q-Q) plots were also performed to assess the overall distribution of p-values and evaluate whether any genomic inflation or deflation occurred though the calculation of lambda (λ) using *QQperm* R package (Petrovski and Wang 2016). All models with genomic inflation (λ) below 0.8 or higher than 1.2 were not considered. Manhattan and Q-Q plots were generated using *CMplot* R package (Lilin-Yin 2023).

### 2.6 Candidate genes and pathways selection

To identify potential candidate genes, we scrutinized the genomic regions surrounding the significant DArTseq markers using the Cameor and ZW6 pea reference genome browsers (https://urgi.versailles.inra.fr/jbrowse/gmod_jbrowse/?data=myData/Pea/Psatv1a/data; https://www.peagdb.com/index/).

Based on the linkage disequilibrium resulted in this pea core collection described by Rispail et al. (2023), we considered genes either containing a DArTseq marker or residing within a 30 kb window from the marker as putative candidate genes.

The process of gene annotation involved associating these candidate genes with enzyme codes to investigate their metabolic pathways using the Kyoto Encyclopaedia of Genes and Genomes (KEGG). In cases where the proteins encoded by these genes were not characterized in the KEGG database, we extended our search to include orthologous sequences. For this purpose, the TAIR database was mined to identify corresponding orthologous proteins in *Arabidopsis thaliana*. This extensive annotation approach facilitated the elucidation of the potential function of each gene and the delineation of pathways relevant to resistance mechanisms against pea rust. To enrich our analysis, we integrated this genomic data with existing literature and performed comprehensive *in silico* analyses, thereby enhancing our understanding of the genetic factors contributing to rust resistance in peas.

## 3 Results

### 3.1 Descriptive statistic, variance components, and principal component analysis

In response to the UpCo-01 isolate, all genotypes exhibited rust symptoms under both controlled conditions (CC) and field conditions, showing a diverse range of trait responses (Table 1). Most accessions demonstrated moderate symptom levels, except for Infection Type (IT), where a predominant presence of well-formed pustules (IT = 4) was observed, as indicated by the skewness for IT (-1.94). The Area Under the Disease Progress Curve for Infection Frequency (AUDPC_IF) and Monocyclic Disease Progress Rate for Infection Frequency (MDPr_IF) both highlighted the disease’s progression with comparably high heritabilities (H^2^ = 0.77 and H^2^ = 0.73, respectively). Despite this, they varied in their descriptive statistics; MDPr_IF showed closer to a normal distribution than AUDPC_IF, as inferred from their skewness. The coefficient of variation also underscored greater disparities between these traits, evident from a higher phenotypic range in AUDPC_IF (173.07) than in MDPr_IF (16.13).

**Table 1.**
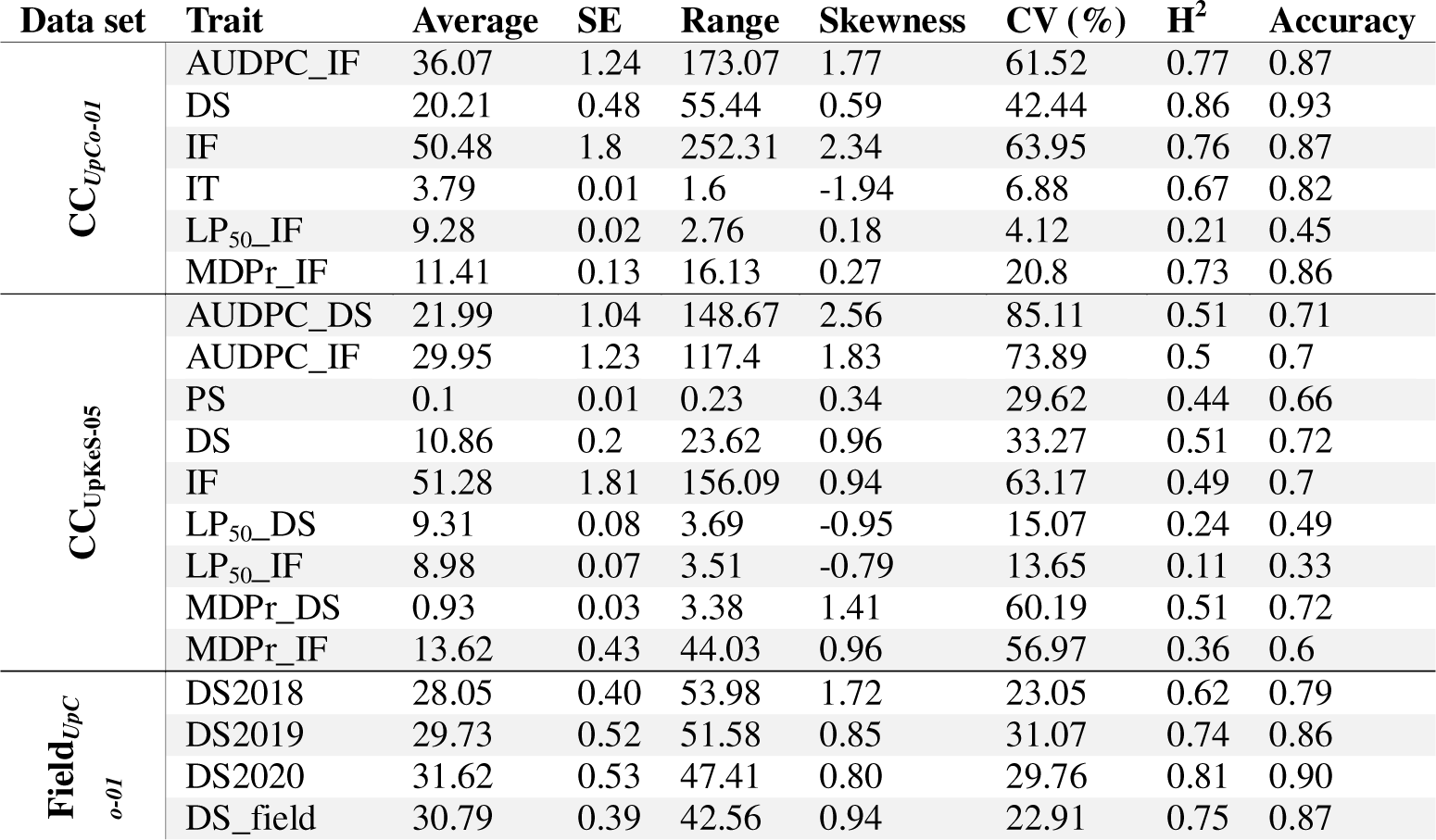
Variance components, heritability and descriptive statistic by phenotypic data set and its trait.

A similar pattern resulted from the evaluation against UpKeS-05, with all accessions displaying rust symptoms (Table 1). In this case the accession PI273209 also displayed necrotic spots reflecting a low IT and a late-acting hypersensitive response to rust (ESM_1). AUDPC exhibited greater skewness and a broader range of values compared to MDPr. For this isolate, however, heritabilities were lower, indicating a distinct virulence pattern. The latency period was the trait with the lowest heritabilities and coefficients of variation for both isolates, not reflecting a solid genetic component behind these variations. Therefore, not considered a selectable trait for breeding perspectives in pea rust.

Field conditions revealed a slightly different picture (Table 1). As a result of the polycyclic evolution of the disease in the field, DS in the three field seasons were higher than in controlled conditions for the same isolate. While the three campaigns and their estimated average (DS_field) had similar mean values and ranges, the 2018 campaign displayed a lower H², accompanied by a pronounced positive skewness of 1.72. This suggests that most pea accessions exhibited reduced DS in 2018 campaign which is further corroborated by its lower CV% relative to the subsequent two field campaigns.

In general, the accuracy values—interpreted as the square root of H²— consistently exceeded 0.7 for most traits, confirming the reliability of the experimental measurements. On the other hand, strong positive (> 0.50) and strong negative (< -0.50) skewness was detected for most traits confirming the need to transform raw values to achieve a normal distribution before fitting the mixed models and predicting the estimated means (BLUEs) for each trait.

The PCA biplot provides a comprehensive visual representation of the primary patterns of variability among the nineteen traits across the three datasets (Fig. 1). The first two principal components (PCs), accounted for over 50% of the total trait variability. Dim1 differentiated traits based on their influence on rust symptoms. Traits positively contributing to rust symptoms clustered on the right, while those inversely related to disease severity, such as latency periods, are positioned on the left although with a lower contribution to the variability. In contrast, Dim2 effectively delineated traits based on the rust isolate inoculated for the evaluations. Specifically, traits assessed for UpCo-01 were situated at the top of the biplot, which combine field and CC_UpCo-01_, while those associated with UpKeS-05 isolate are anchored at the bottom.

**Fig. 1.**
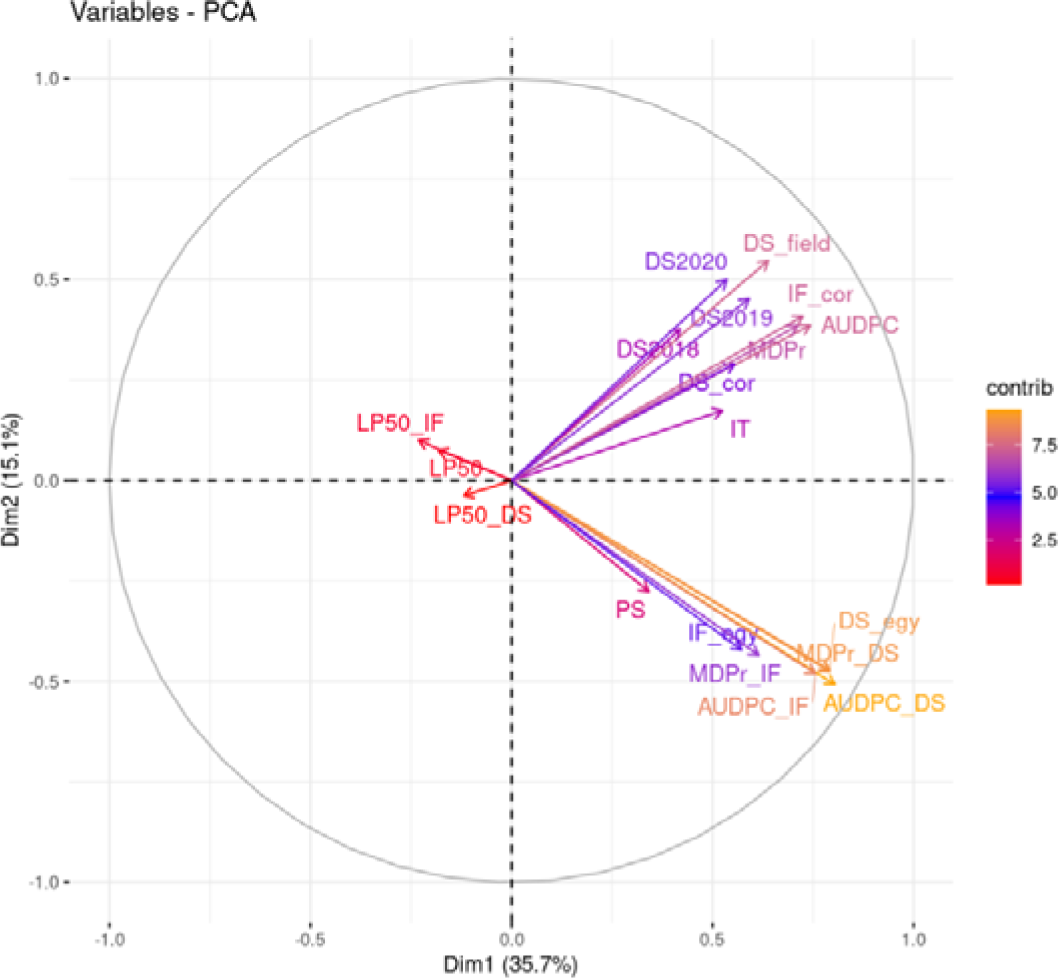
PCA biplot by trait contribution to phenotypic variance.

### 3.2 Detection of Associated Markers

To identify genetic variations associated with rust symptoms in pea, a GWAS was conducted using two different models, MLM and BLINK, for each of the 19 traits evaluated in three conditions across 320 pea accessions. A total of 95 DArTseq markers were found to be significantly associated with some of the evaluated traits in combination with the two models applied (Fig. 2 and ESM_2). Examining the BIC estimates, the genomic inflation factor (λ) and the QQ plot revealed that population structure was efficiently controlled by the Kinship matrix without the need to add other population structure covariates (PCs) (Fig. 3, Table 2, and ESM_3).

**Fig. 2.**
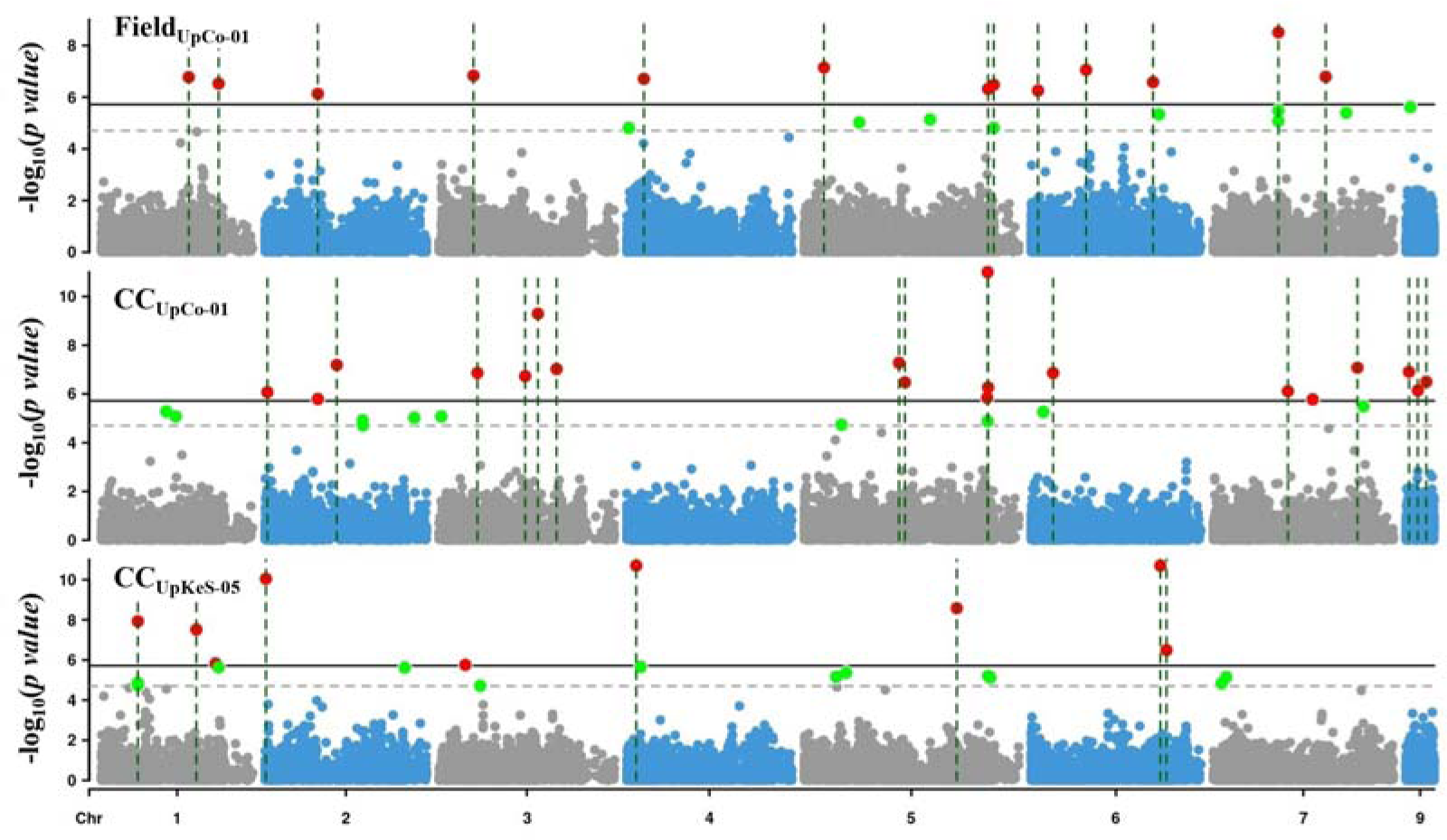
Manhattan plots showing marker significance in a combination of MLM and BLINK models by phenotypic data base across pea genome. Chromosome 9 shows unmapped markers onto reference genome. Associated markers through Bonferroni-based LOD are highlighted in red and the genomic region with dashed lines while in green are the associated markers retrieved based on FDR adjustment method.

**Fig. 3.**
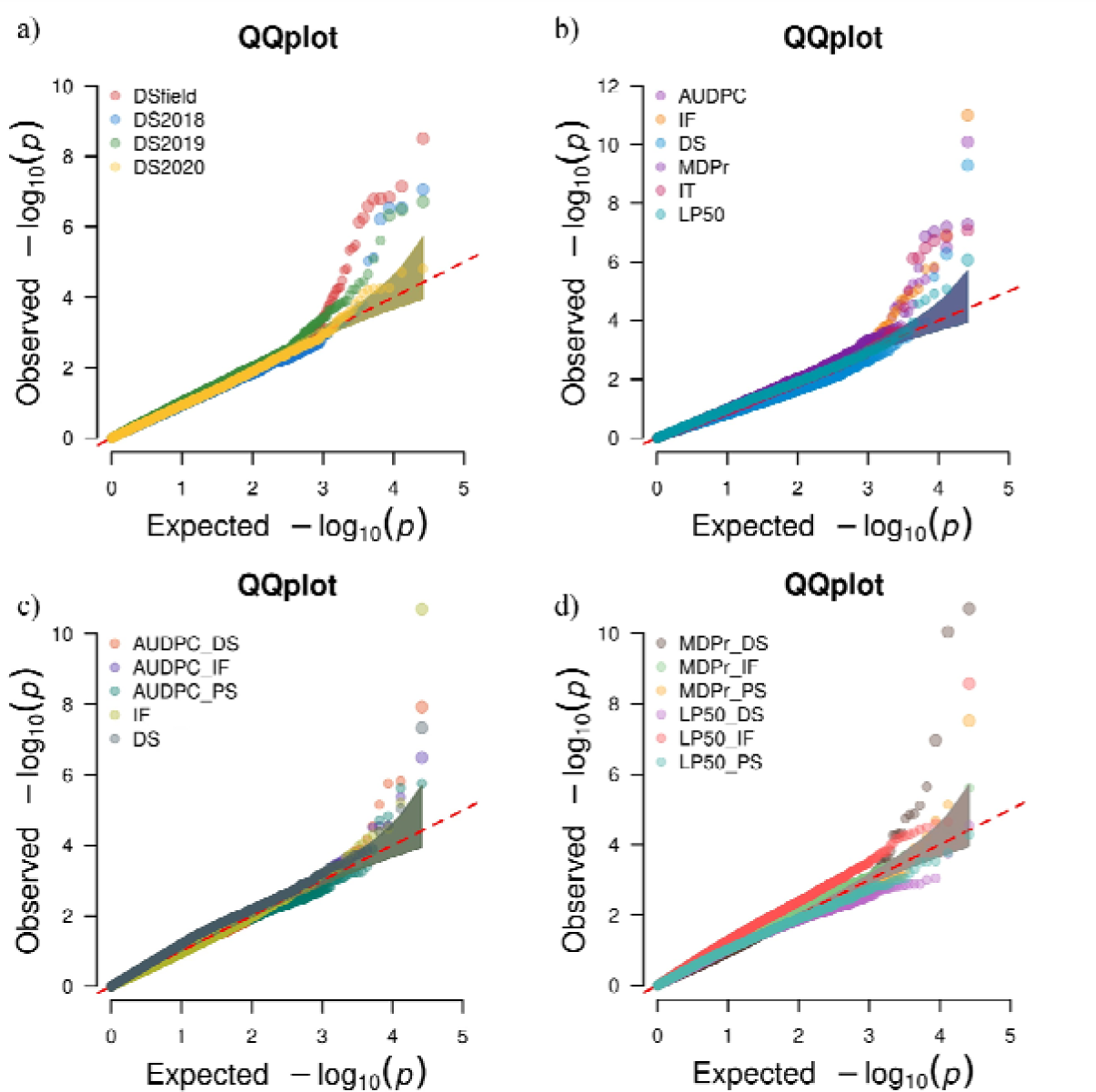
Q-Q plots between theoretic and observed p values using the BLINK model. a) and b) show p values from UpCo-01 isolate in field and controlled conditions, respectively. c) and d) show p values from UpKeS-05 results in controlled conditions.

**Table 2.**
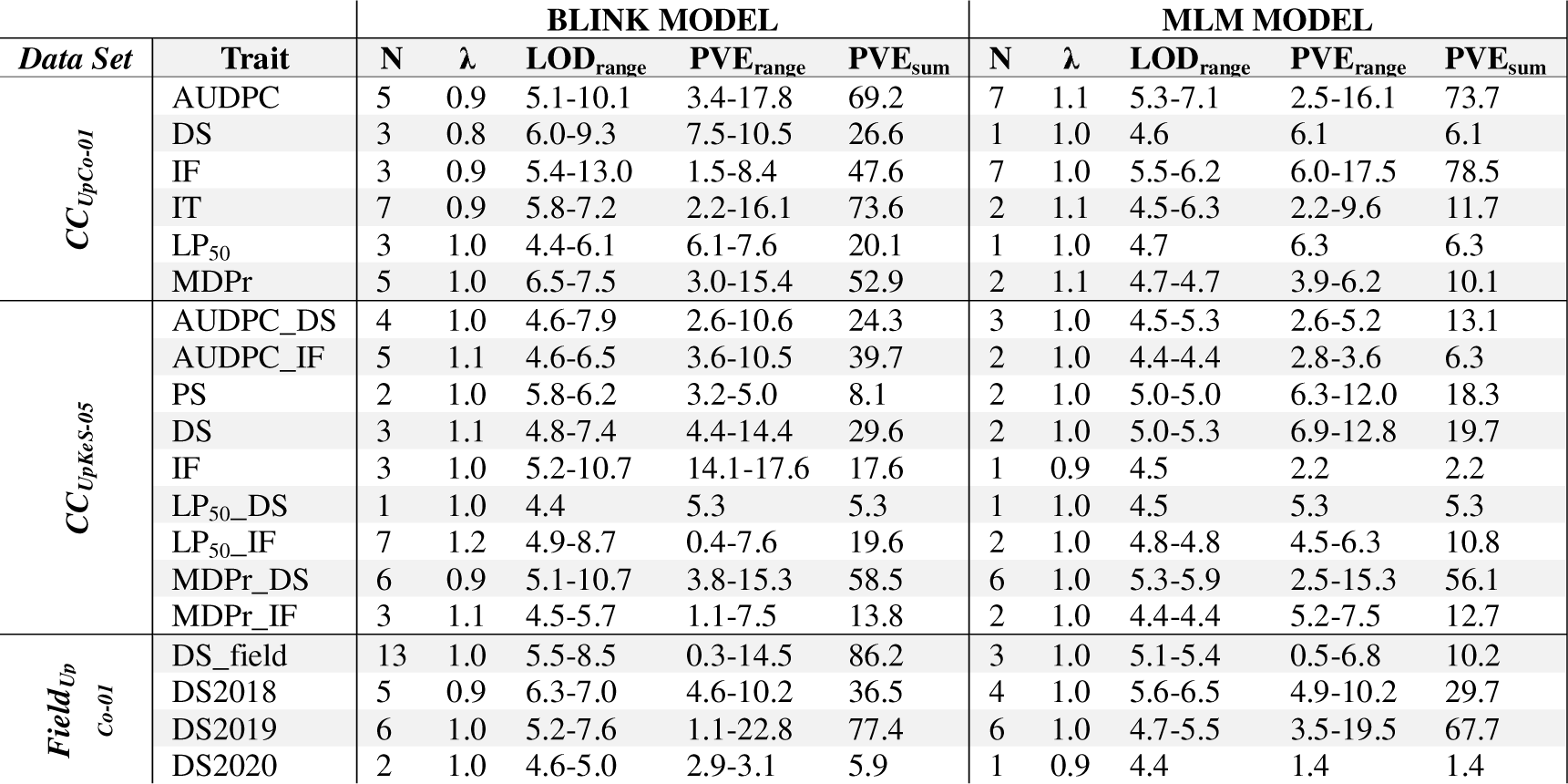
BLINK and MLM model outputs obtained for each trait from the three phenotypic data sets. N indicates the number of significant marker-trait associations; λ is the genomic inflation factor; the LOD_range_ is the range of the -log_10_(p-value); PVE_range_ is the range of the phenotypic variance explained by each individual marker and PVE_sum_ is the total phenotypic variance explained by the associated markers.

The BLINK approach identified 70 markers linked to pea rust of which 49 were unique to this model while the MLM model detected 46 associated markers of which 25 were unique to this method. Notably, 21 associated markers were common to more than one trait or model (Fig. 4a) while one marker was identified from all three data sets (Fig. 4b).

**Fig. 4.**
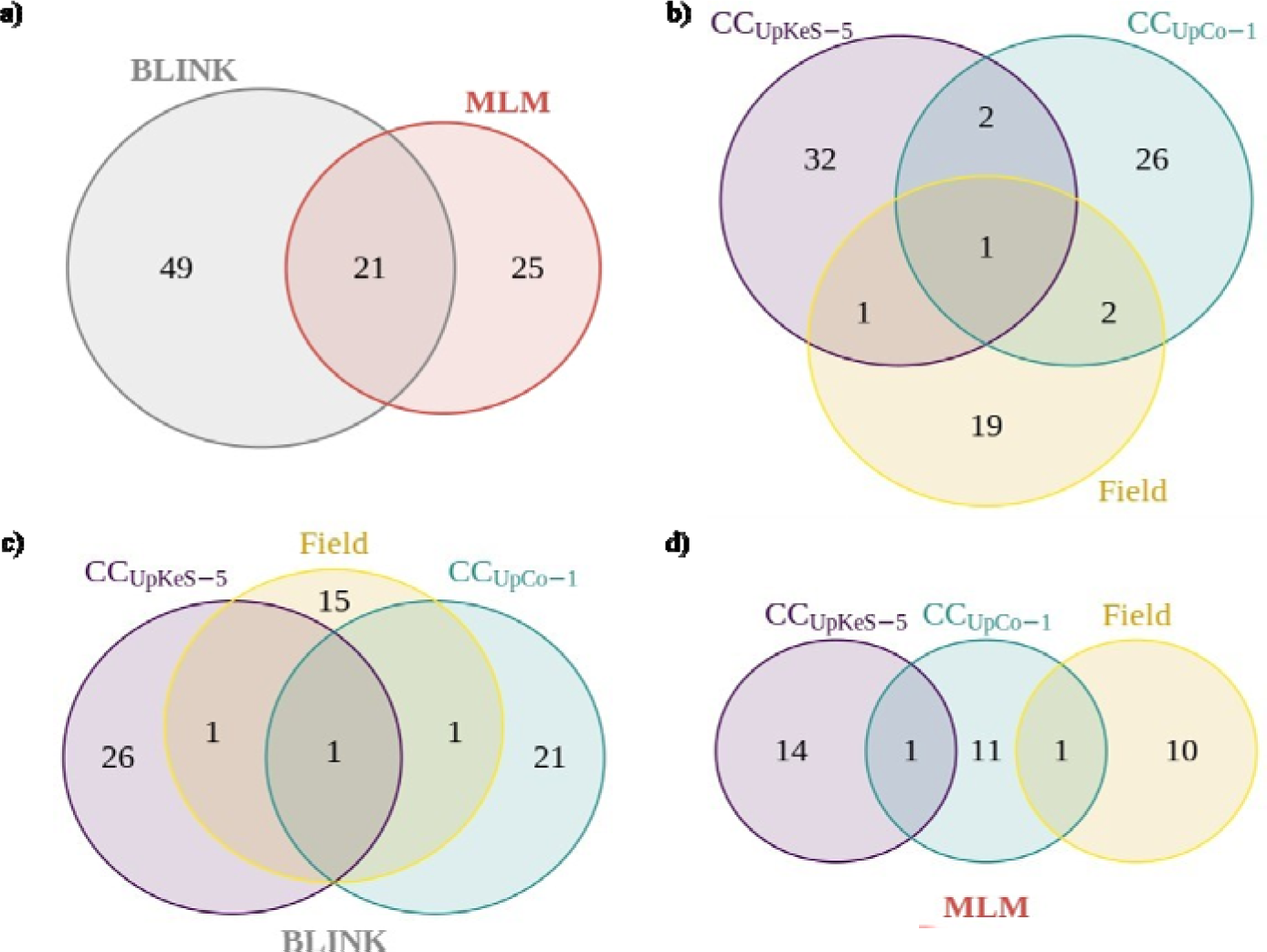
Venn diagram of common markers across data sets. a) and b) represent the common associated markers by applied model and the common associated markers by phenotypic dataset in combination of both models’ outputs. c) and d) show the linked markers to pea rust from BLINK and from MLM results, respectively.

In general, the BLINK model yielded a higher number of significantly associated markers with lower p-value and higher LOD (Table 2). The total phenotypic variation explained by the associated marker for each trait (PVE) varied also greatly depending on the GWAS model. As the BLINK model identified more significant marker-trait associations for each trait, their corresponding PVE_sum_ was generally higher (Table 2). The highest number of associated markers for a single trait was identified with the BLINK method for DS_field, with a total of 13 markers explaining 86.2% of the total phenotypic variance while the MLM model only uncover 3 associated markers for this trait accounting for 10.2% of the phenotypic variance (Table 2). By contrast, the MLM method uncovered 7 markers associated with AUDPC for CC_UpCo-01_ while BLINK only uncovered 5 significantly associated marker for this trait although associated markers identified by both methods explained around 70% of PVE_sum_.

It is worth noting that the sum of the phenotypic variation explained by the associated markers for each trait was closely related to heritability for the BLINK model (ESM_4), except for DS2020.

### 3.3 *In silico* identification of candidate genes

To identify putative candidate genes the genomic regions surrounding the significantly associated markers was examined. Out of the 95 associated markers detected (ESM_2), 75 markers were located within or next to 62 genes from the Cameor and/or ZW6 reference genomes, referred in Table 3 by their NCBI gene ID codes. It is interesting to note several of these candidate genes contained more than one significant associated marker, underscoring their importance in contributing to the phenotypic variation of the trait. By chromosome, the gene annotation results were as follows:

**Table 3.**
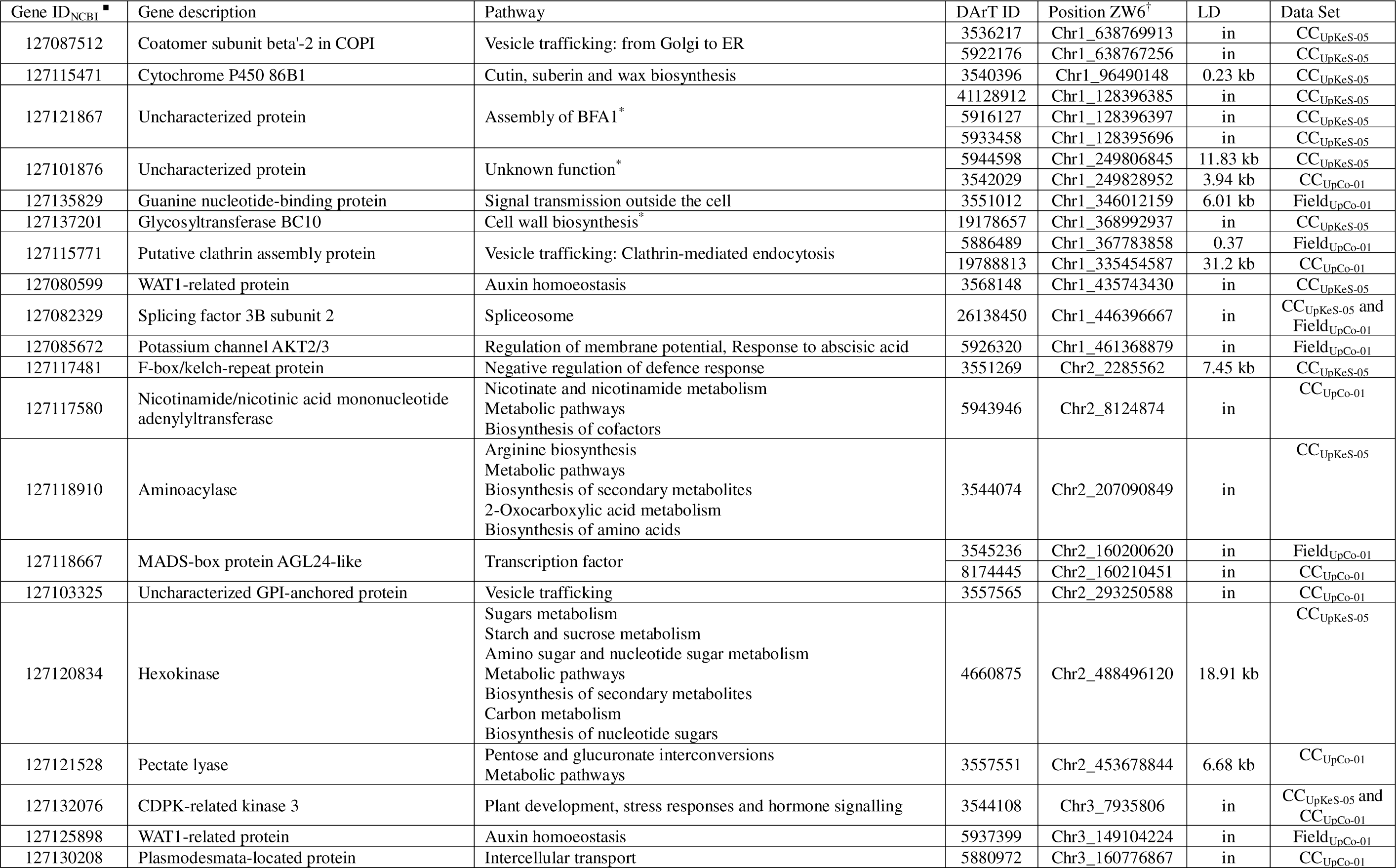
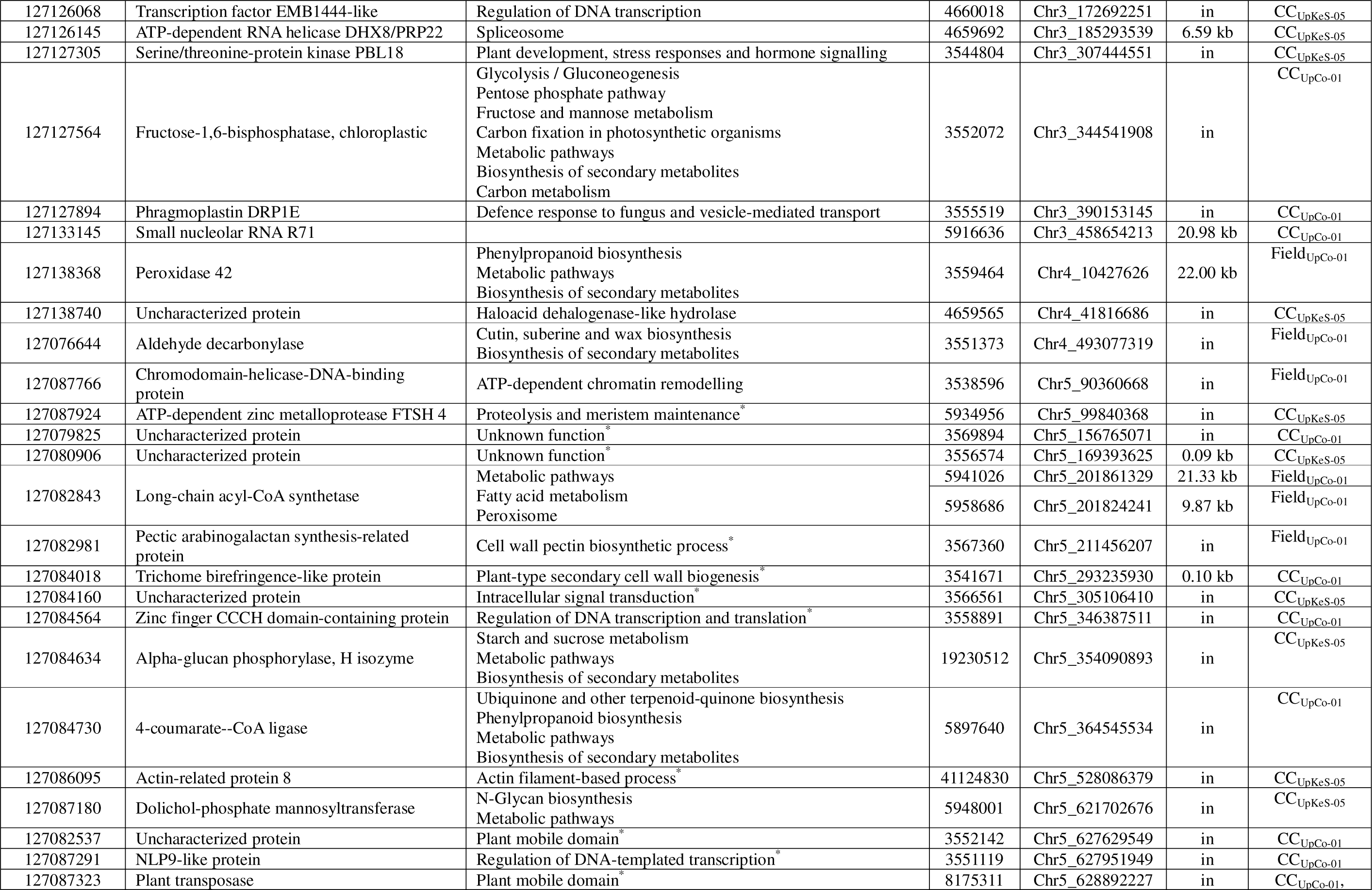
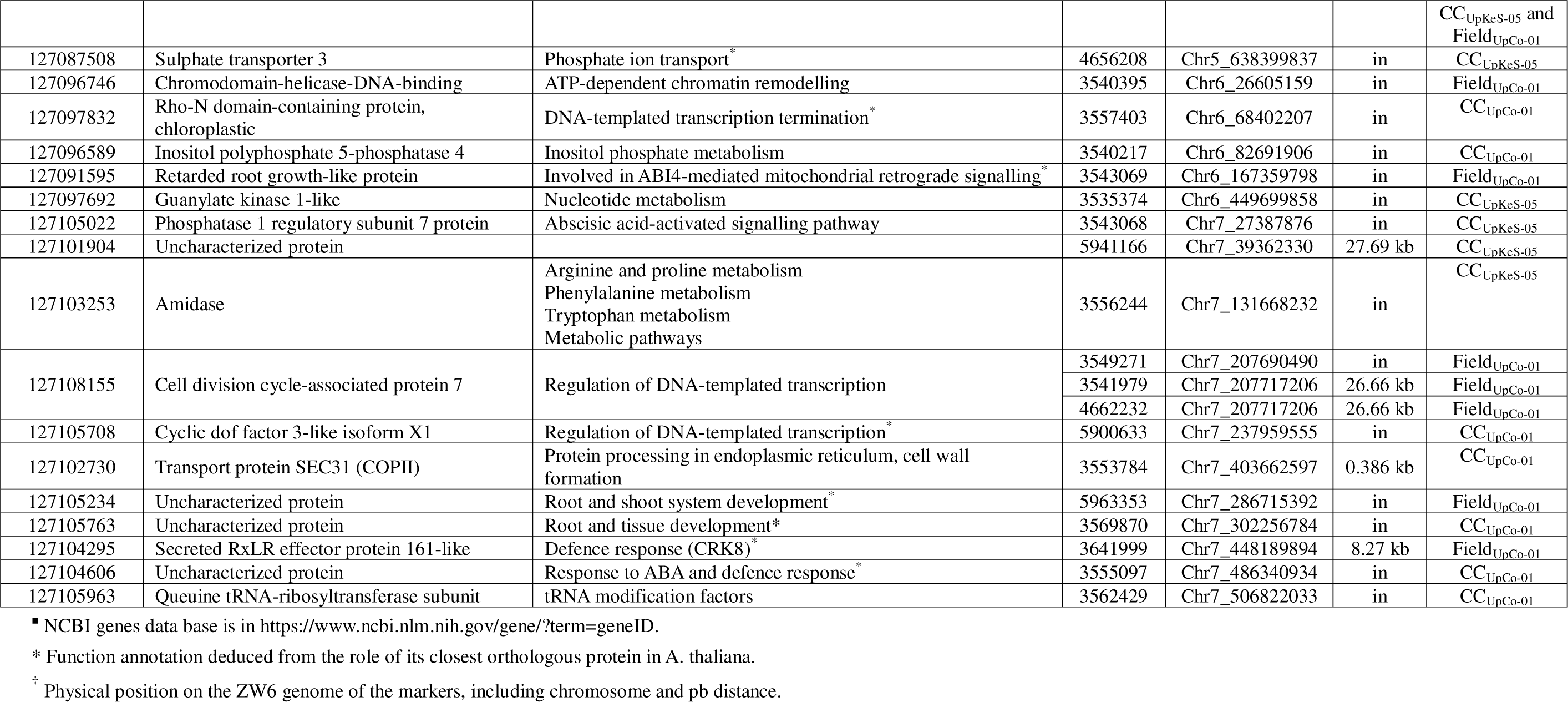
Potential candidate genes containing or in the vicinity of significant markers.

A total of 10 candidate genes were detected on chromosome 1. Among them, two genes, 127087512 and 127115771, each marked by two associated markers were related to vesicle trafficking within cells. Two additional genes 127115471 and 127137201 were related with wax and cell wall biosynthesis. Interestingly, one uncharacterized gene, marked by three associated markers, was related to BFA1 assembly which is required for ATP synthesis in *A. thaliana*.

On chromosome 2, 7 candidate genes were detected. Four of these genes (127120834, 127121528, 127118910 and 127117580) are related with carbohydrate and amino acid metabolism. Two genes, 127117481 encoding an F-box protein, and 127118667 encoding a MADS-box transcription factor were related to defence response to stresses in plants (Table 3) while the last identified candidate genes engaged in cell trafficking.

On chromosome 3, 9 candidate genes were detected. Among them, two genes, 127132076 and 127127305, encoded protein kinases related to plant development, stress responses and hormone signalling. Two genes, 127130208 and 127127894, were also associated with vesicle trafficking. Additionally, two genes were related with secondary metabolism while another one was related to auxin homeostasis as gene 127080599 from chromosome 1.

From the three genes detected on chromosome 4, two genes (127138368 and 127076644) encoded proteins related with the secondary metabolism and cell wall biosynthesis while the last one, gene 127138740 putatively encodes a haloacid dehalogenase-like hydrolase (HAD).

Chromosome 5 harboured the highest number of candidate gene with 18 genes identified from 23 associated markers. A notable hotspot in chromosome 5 consisted of three closely spaced markers within a 1.2 Mb window, located within genes 127082537, 127087291, and 127087323. Interestingly, two of these genes 127082537 and 127087323 were related with mobile elements in *A. thaliana* while the third one together with genes 127087766 and 127084564 were related with DNA transcription regulation. As in chromosomes 1 and 4, several genes related with cell wall biosynthesis and maintenance were detected on chromosome 5. Several genes related with transport and signalling, and with fatty acid or secondary metabolism were also detected on this chromosome.

From the five candidate genes identified on chromosome 6, three were related with DNA formation and transcription. The last two genes were related to inositol phosphate metabolism and abscisic acid (ABA) signalling, respectively.

Chromosome 7 harbour 11 genes in the vicinity of 13 associated markers. Among them, the function of three genes were not known in peas but the ortholog of two of them, 127105234 and 127105763, in *A. thaliana* have been related to tissue development. Similarly, to the other chromosomes, several candidate genes related with the regulation of DNA transcription regulation (127108155 and 127105708), the amino acid metabolism (127103253), the vesicle trafficking (127102730) and the response to stresses (127105022, 127104606 and 127104295) were identified in this chromosome.

## 4 Discussion

Rust is a world-wide distributed disease that severely affects pea crop yield (Rubiales et al. 2015). In warm and humid areas, the main causative pathogen is *Uromyces viciae-fabae*, while *U. pisi* is responsible for pea rust in more temperate regions (Singh et al. 2023). Complete resistance to rust in peas has not yet been identified, with partial resistance being the primary source of genetic resistance available so far to control this disease in pea (Barilli et al. 2014). In this study, we applied a GWAS approach to identify regions of the pea genome associated with traits linked to the partial resistance response. To achieve this, a collection of 320 pea accessions was evaluated under field and control conditions in response to two rust isolates (UpCo-01 and UpKeS-05).

### 4.1 Phenotypic variance and GWAS model outputs

The evaluation method and phenotypic response of the collection against the rust isolate UpCo-01 was previously described in Osuna-Caballero et al. (2022). Here, we describe the phenotypic response of the same collection against the rust isolate UpKeS-05 obtained using the methodology described in Osuna-Caballero et al. (2023). While the seedlings inoculation followed the same procedure, this evaluation performed by images using the R software for the UpKeS-05 isolate in detached pea leaflet revealed a concordance correlation coefficient between the visual evaluation of 0.962 according to Osuna-Caballero et al. (2023). Therefore, in this study the effect of the isolates can be compared despite having obtained the phenotypic data in two different ways: an evaluation by visual counting (Osuna-Caballero et al. 2022) and another evaluation based on images using R software (Osuna-Caballero et al. 2023). These results comparison revealed some commonalities and differences. Similarly, to the situation with UpCo-01, complete resistance to UpKeS-05 was not detected in the collection. In addition, the reduced number of well-developed pustules and low IT detected on the accession PI273209 in response to UpKeS-05 suggest that this accession also exhibits a late hypersensitive resistance response against this isolate (ESM_1). Disease ratings were nonetheless lower in response to UpKeS-05 than to UpCo-01 and the response to each isolate were clearly differentiated on the PCA suggesting that the collection responded differentially to the isolates. As mild to moderate relationship was also observed for most traits between isolates, these differences are more likely revealing differences in the overall virulence level of the isolates although the existence of *U. pisi* races in peas could not be ruled out.

Previous studies performed on the pea core collection demonstrated a rapid LD decay and a broad distribution of the polymorphic DArT markers throughout the pea genome (Rispail et al. 2023). Although a significant portion of the population resulted admixed in phylogenetic terms, the collection was found to be structured into six groups consistent with the taxonomy of the *Pisum* species and subspecies (Rispail et al. 2023). However, BIC selection models indicated that population structure was efficiently controlled in the GWAS models by the kinship matrix alone and did not require adding the initial PCs to the models (ESM_3). This was previously demonstrated on GWAS studies performed on plant populations with a high percentage of admixture (Kaur et al. 2023). The BLINK model improves the statistical power and enhances the robustness of association signals compared to MLM (Huang et al., 2019). Furthermore, the extensive coverage of the DArT markers across the pea genome is ideal for multi-locus methods like BLINK, making it a frequently used model in GWAS studies to analyse complex traits in legumes (Tibbs Cortes et al. 2021; Susmitha et al. 2023). Accordingly, this method detected a high number of markers associated with rust disease development.

The two GWAS models applied here detected a total of 95 markers significantly associated with one or more of the 19 rust disease traits, among which 21 markers were detected by both methods which will be instrumental for future breeding for rust resistance in pea. The current genetic sources of pea resistance against *U. pisi* are described in the work of Barilli et al. (2010,2018). Three QTLs have been described to date. The Up1 QTL identified in linkage group 3 (chromosome 5) by Barilli et al. 2010 explained a total phenotypic variance of 63% in the biparental F_2_ population between *P. fulvum* (resistant) and *P. sativum* (susceptible). These results have not been corroborated in this work, where the markers on chromosome 5 are not close to the RAPD markers that flank the QTL Up1: OPY11_1316_ and OPV17_1078_. The subsequent results of Barilli et al. 2018 also did not corroborate Up1 but two new QTLs UpDSII and UpDSIV were identified in linkage groups 2 and 4, corresponding to chromosomes 6 and 4, respectively. The QTL UpDSII explained an average of 24% of the phenotypic variation in the field and 28% under controlled conditions. Similarly to our results, the markers found around UpDSII revealed 16% phenotypic variation in the field and 20% for the same UpCo-01 isolate. On the other hand, the QTL UpDSIV explained 20% of the phenotypic variation both in the field and under controlled conditions according to the results of Barilli et al. (2018). Markers colocalized near UpDSIV in this work revealed an explained phenotypic variation of 26% in the field. However, no markers associated with the severity of rust caused by *U.pisi* (UpCo-01) were found on chromosome 4. These discrepancies may be due to the origin of the biparental population, due to interspecific crossing between *P. fulvum* and *P. sativum*, in counterpart of the GWAS population carrying greater genetic diversity. On the other hand, this is the first GWAS targeting rust resistance in pea while similar studies addressing rust disease in legumes remains scarce. Most previous studies were based on the MLM method while only one study on soybean (*Glycine max*) – rust (*Phakopsora pachyrhizi*) applied the BLINK method (Martins et al. 2022; Montejo Domínguez et al. 2022; Wu et al., 2022; Xiong et al. 2023). The number of significant marker-trait associated largely depended on the number and distribution of the available markers. Accordingly, 100 and 129 marker-trait associations were identified in response to rust in soybean and common bean in studies with more than 20,000 genome-wide markers (Wu et al. 2022; Xiong et al. 2023) while only 7 DS associated markers was identified in response to *U. pisi* in grass pea from a marker set of 5,651 SNPs (Martins et al. 2022). Interestingly, aligning these 7 grass pea markers onto pea reference genomes allowed the identification of 19 candidates genes among which 5 have similar function to those detected in the present study (Martins et al. 2022).

### 4.2 Candidate genes represent diverse functional roles

Examining the surrounding regions of the 95 associated markers detected in the present study allowed the identification of 62 candidate genes with potential function in rust resistance. According to their annotation, these candidate genes participate in a variety of function, including primary and secondary metabolism, cell wall synthesis, cell trafficking, DNA transcription regulation and defence response.

#### 4.2.1 Regulators of gene expression

Among the 62 candidate genes, 12 are related to the regulation of DNA transcription/translation, RNA modification, and transposable elements. Seven of them are transcription factors (TFs), including a Nodule inception-like 9 protein (NLP9, 127087291) a Cycling DOF factor 3 (CDF3, 127105708) and a Zinc finger transcription factor orthologous to Tandem Zinc Finger 9 (TZF9, 127084564). Previous studies on TZF9 identified it as an important regulator of plant immune defence in *A. thaliana* (Maldonado-Bonilla et al. 2014; Tabassum et al. 2020). While NLP9 and CDF3 have been mainly associated with nitrogen use efficiency in plants some studies in tomato and soybean linked them to the response to biotic and abiotic stresses (Renau-Morata et al. 2017; Domínguez-Figueroa et al. 2020; Konishi et al. 2021; Amin et al. 2023). Interestingly, two candidate genes (127087323 and 127082537) encoding plant mobile domains are located within a 1.2Mb distance on both side of NPL9. While the role of these two genes in genomic regulation is under study, their involvement in plant defence has yet to be clarified (Fambrini et al. 2020). Apart from these TFs, two candidate genes (12782329 and 127126145), encoding a splicing factor (SF3b) of the U2 spliceosome complex and an RNA helicase (PRP22) respectively, are related to mRNA splicing via the spliceosome, which play an important role in the plant response to biotic stresses (Shang et al. 2017). Accordingly, knock-out mutation in SMD3, other subunits of the U2 complex, increase the susceptibility to *Pseudomonas syringae* in *A. thaliana* (Golisz et al. 2021). In addition, the orthologous gene of RNA helicase in *A. thaliana* (At1g32490) have been implicated in the negative regulation of numerous genes related to cell wall formation (Howles et al. 2016), which could improve the physical barrier against haustoria formation inside the plant cells.

#### 4.2.2 Regulators of vesicle trafficking and cell wall components

Several of the candidate genes containing or closely related to the associated markers are related to vesicle trafficking and cell wall biosynthesis suggesting the importance of the cell wall and cell trafficking in response to rust. Among them, three genes (127087512, 127102730, and 127115771) encode proteins of the coat protein complex I (COPI), coat protein complex II (COPII), and clathrin-mediated endocytosis vesicles that mediate basic functions of protein export/import to the ER in plant cells (Zeng et al. 2023). COPII has been directly involved in the process of autophagy, which enable the ABA-mediated cell nutrient recycling in response to stress (Li et al. 2022). In addition, the gene 127086095 encodes an actin-related protein (ARP8) that participates in cytoskeleton-related functions (Kandasamy et al. 2008). Recent studies in *A. thaliana* have shown that related ARPs, such as ARP4 and ARP6, are involved in both biotic and abiotic environmental changes (Nie and Wang 2021; Jakada et al. 2023). This suggest that vesicle trafficking, and particularly the endomembrane cell system, could play an important function in response to rust in pea.

On the other hand, eight candidate genes (127086095, 127076644, 127115471, 127137201, 127084018, 127082981, 127076644 and 127115471) are related to cell wall formation or to the biosynthesis of cell-wall related secondary metabolites such as lignin. Among them, gene 127137201 encodes a glycosyltransferase BC10 that have been related to the cellulose and lignin biosynthesis at the cell wall (Zhou et al. 2009; Zhang et al. 2016) while two genes (127082981 and 127084018) are related to the synthesis of rhamnogalacturonan-I (RG-I), one of the pectic components of the cell wall (Stonebloom et al. 2016). This suggests that changes in the cell wall composition may participate in the plant cell response to rust. Interestingly, two additional genes (127076644 and 127115471), encoded an Eceriferum (CER1) -like enzyme and a mid-chain alkane hydroxylase (MAH1), respectively, are important players of wax on the cell wall (Lewandowska et al. 2020; Wang et al. 2020) which play an important role in the host recognition by rust in several species (Niks and Rubiales 2002) and on the modulation of plant susceptibility to necrotrophic fungus such as *Sclerotinia sclerotiorum* (Bourdenx et al. 2011).

### 4.2.3 Regulators of hormone signaling and defense response

Eleven of the identified genes in the vicinity of the significantly associated markers were related to hormone signalling and defence. Among them, two genes (127080599 and 127125898) encoded two copies of a nodulin like proteins homologs to *Medicago truncatula* NODULIN 21 (MtN21) and *A. thaliana* WAT1 (walls are thin 1) genes that have been involved in auxin metabolism and play a significant role in secondary cell wall formation of (Ranocha et al. 2010). Recent studies showed that additional MtN21-related homologs in *A. thaliana*, such as RTP1 negatively regulates plant resistance to biotrophic pathogens through cell death and auxin-mediated production of reactive oxygen species (Pan et al. 2016; Gao et al. 2023) linking these genes to auxin signalling and defence. Apart for auxin, several genes pointed to the potential involvement of ABA in response to rust since four genes (127085672, 127091495, 127105022 and 127104606) encoded ABA-responsive genes including orthologs of the potassium channel AKT2/3, the regulatory subunit of protein phosphatase 1 (PP1R3), the Abscisic Acid Insensitive 4 (ABI4) and a SEC14-like protein (Chérel et al. 2002; Wang et al. 2016; Chandrasekaran et al. 2020; Zhang et al. 2020). Among their distinct functions, several studies in *A. thaliana* or *Nicotiana benthamiana* showed that AKT2/3, ABI4 and the SEC14 proteins contributed to plant defence being important actors of the plant resistance to different pathogens (Kiba et al. 2012; Zhou et al. 2014; Yop et al. 2023).

Several genes identified in this study are also directly involved in the plant defence mechanisms against pathogen attacks, involving some regulatory proteins and kinases. Accordingly, three of the identified genes encoded protein kinases including a PBS1-like protein kinase, a calcium-dependent protein kinase (CDPK3) and cysteine-rich receptor-like kinase (CRK8). Previous studies reported the crucial role of PBS1 in bacterial and viral resistance in *A. thaliana* and soybean (Pottinger et al. 2020). PBS1 was also shown to play a role in herbivore resistance in interaction with CDPK3 (Miyamoto et al. 2019; Desaki et al. 2023) hinting the involvement of both gene in a common defence pathway. Interestingly, one marker significantly associated with pea rust resistance was located at 8.27 kb of the receptor-like kinase CRK8 that have been shown previously to play a crucial role in rust resistance in wheat (Gu et al. 2020; Kamel et al. 2023). In addition to these kinases, this study identified an ortholog of a F-box protein CPR1/CPR30 (127117481), that acts, between other function, as a negative regulator of salicylic acid-dependent resistance (R) proteins like SNC1 (Gou et al. 2012), and a dynamin-related protein 1E (DRP1E; 127127894) which knock-out mutation was found in *A. thaliana* to induce stronger hypersensitive response to powdery mildew (Tang et al. 2006; Leibman-Markus et al. 2022; Mc Gowan et al. 2022).

Altogether, this GWAS study shed light on the complex genetic architecture underpinning partial resistance to rust in pea. In addition to the detection of 95 new molecular markers associated with partial resistance, this study detected 62 candidate genes that contains or was intricately linked to these markers, which offer several promising targets to improve our understanding of rust resistance in pea and future plant breeding strategies. These genes span a diverse array of biological functions providing the basis for further gene functional mapping. The identification of several candidate genes known to participate in rust resistance in cereal (i.e., wax genes, CRK8) suggest that at least some of these candidate genes might be the responsible gene underlying the detected QTL confirming the usefulness of GWAS studies to uncover new resistance QTL and genes. Therefore, this study opens the possibility to characterize further, via gene expression studies or TILLING, the function of these candidate genes to explore their implication in rust disease in pea. The significantly associated DArT-seq markers also provides a basis for future marker-assisted selection and the development of more efficient genomic prediction models for rust resistance. Therefore, this study contributes to understanding the genetic make-up of rust resistance in pea and promotes future crop improvement. By focusing on the candidate genes uncovered, breeding programs can develop more targeted approaches, upon their validation, leveraging genetic variation to improve the resilience of pea crops against rust disease. These findings open new avenues for precision breeding, offering novel targets for rust resistance improvement and enhancement of crop resilience and productivity through the development of new pea rust-resistant varieties. This not only contributes to secure yield and quality in the face of biotic stress but also supports sustainable agricultural practices by reducing the reliance on chemical fungicides.

## Supporting information

ESM_1

ESM_2

ESM_3

ESM_4

## 5 Data availability

The DArTseq marker datasets analysed during the current study are available in the Zenodo repository, https://zenodo.org/records/7180467. The phenotypic datasets generated and analysed are available in the GitHub repository, https://github.com/SalvaOsuna/Rust-collection.git.

## 7 Acknowledgements

The authors acknowledge Manuel A. Jiménez-Vaquero for assisting with the pea evaluations under controlled condition with the rust UpKeS-05 isolate.

## 9 Author Contribution Statement

DR and NR conceived the original idea for this study. SOC conducted the experiments and took the lead in manuscript writing. SOC analysed the data with help from NR. SOC designed the figures and tables with input from NR and DR, who also significantly contributed to data interpretation and provided critical feedback that shaped the final version of the manuscript. DR acquired the funding for this project. All authors approved the submitted version.

## 11 Ethics approval

All experiments described in this manuscript comply with the current laws of the country in which they were performed.

