## Supplementary material for "Genome-Wide association study uncovers pea candidate genes and pathways involved in rust resistance": ESM_1

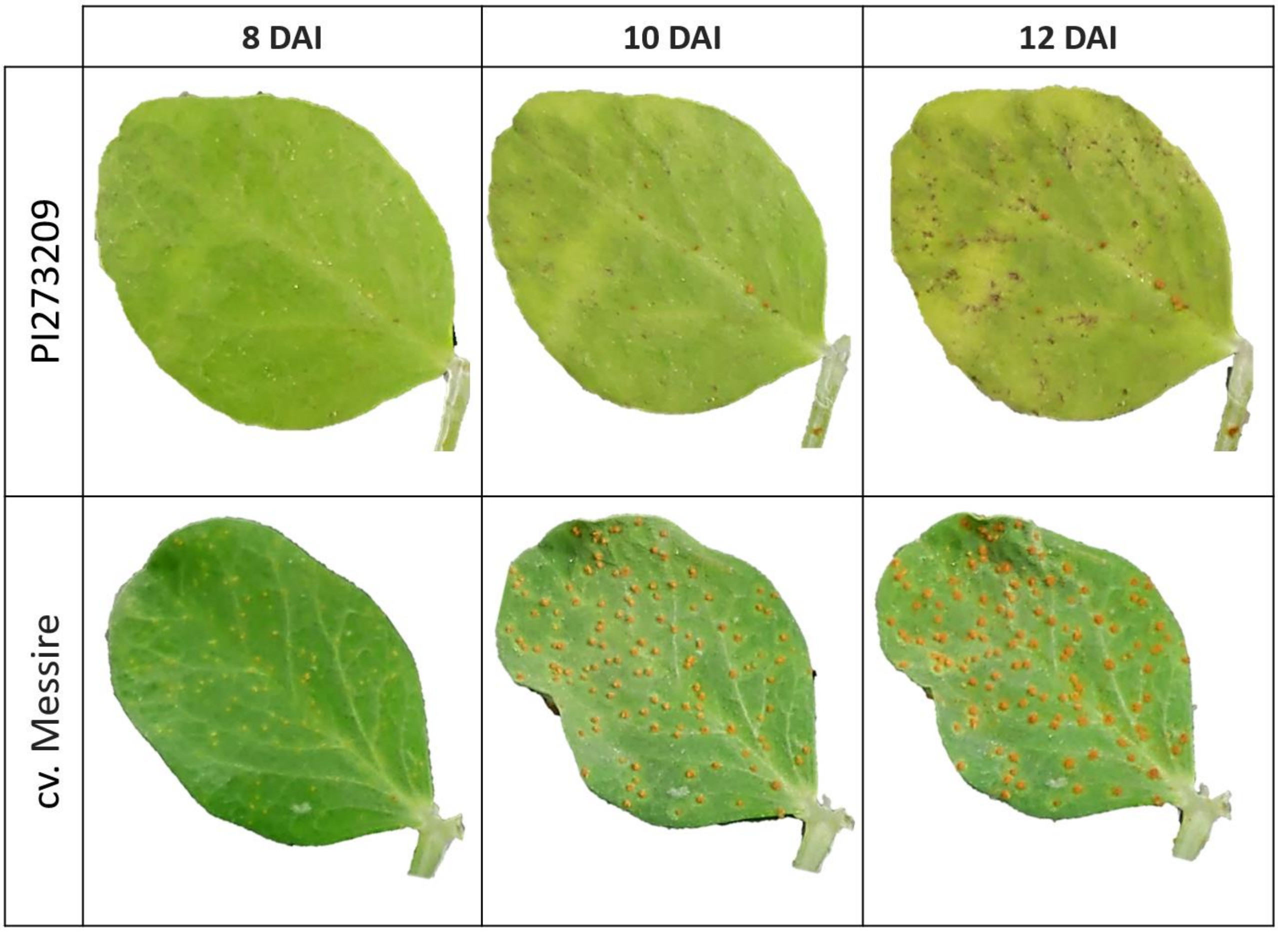


*Figure 1. Rust (UpKe-05 isolate) symptoms progression by 8, 10 and 12 days after inoculation (DAI) in cv. Messire and PI273209 accession leaflets.*
