## Supplementary material for "Genome-Wide association study uncovers pea candidate genes and pathways involved in rust resistance": ESM_4

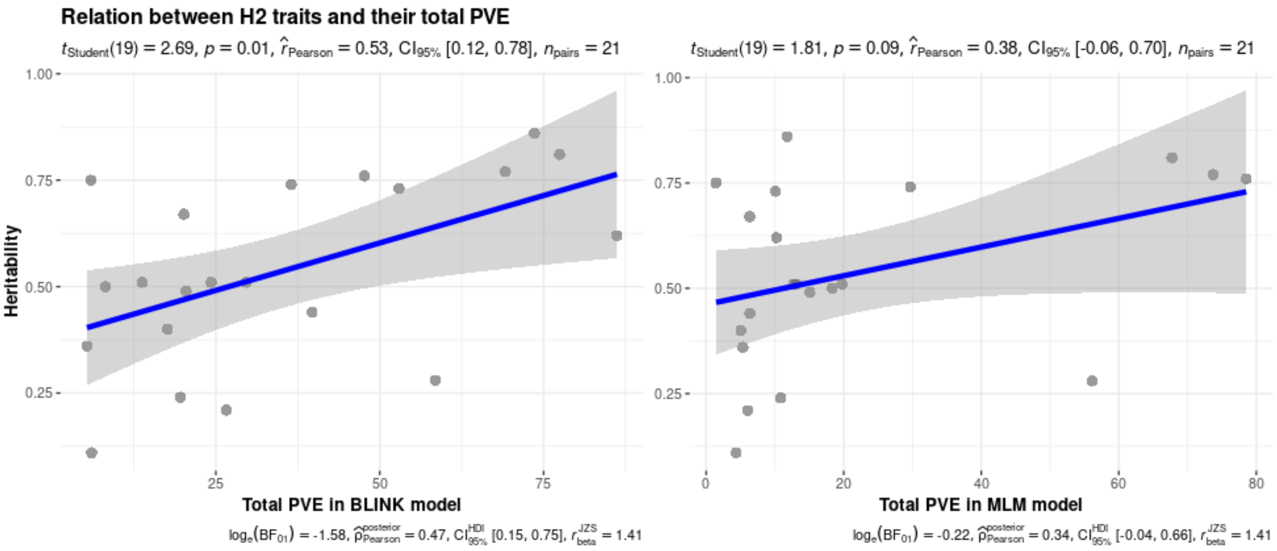


*Figure 1. Scatterplot between heritability of each trait by its cumulative phenotypic variance explained ($PVE_{sum}$) by the markers obtained through BLINK model (left) or MLM model (right). Inferential statistics with effect size plus CIs are at the top of the plot while Bayesian hypothesis-testing and estimation are at the bottom.*
